# Gut Microbe-Targeted Choline Trimethylamine Lyase Inhibition Improves Obesity Via Rewiring of Host Circadian Rhythms

**DOI:** 10.1101/2020.12.04.411546

**Authors:** Rebecca C. Schugar, Christy M. Gliniak, Robert N. Helsley, Amanda L. Brown, Amy Burrows, Chelsea Finney, Kevin K. Fung, Frederick M. Allen, Daniel Ferguson, Anthony D. Gromovsky, Chase Neumann, Amy McMillan, Jennifer A. Buffa, James T. Anderson, Margarete Mehrabian, Maryam Goudzari, Belinda Willard, Tytus D. Mak, Andrew R. Armstrong, Garth Swanson, Ali Keshavarzian, Jose Carlos Garcia-Garcia, Zeneng Wang, Aldons J. Lusis, Stanley L. Hazen, J. Mark Brown

**Affiliations:** Department of Cardiovascular and Metabolic Sciences, Lerner Research Institute Cleveland Clinic, Cleveland, OH 44195, USA; Center for Microbiome and Human Health, Lerner Research Institute, Cleveland Clinic, Cleveland, OH 44195, USA; Department of Cardiovascular Medicine, Heart and Vascular Institute, Cleveland Clinic, Cleveland, OH 44195 USA; Proteomics and Metabolomics Core, Lerner Research Institute, Cleveland Clinic, Cleveland, OH 44195; Departments of Medicine, Microbiology, and Human Genetics, University of California Los Angeles, Los Angeles, CA 90095, USA; Life Sciences Transformative Platform Technologies, Procter & Gamble, Cincinnati, OH, USA; Department of Internal Medicine, Division of Gastroenterology, Rush University Medical Center, Chicago, IL 60612, USA; Mass Spectrometry Data Center, National Institute of Standards and Technology (NIST), Gaithersburg, MD 20899, USA

**Keywords:** obesity, microbiome, insulin resistance, circadian

## Abstract

Obesity has repeatedly been linked to reorganization of the gut microbiome, yet to this point obesity therapeutics have been targeted exclusively toward the human host. Here we show that gut microbe-targeted inhibition of the trimethylamine N-oxide (TMAO) pathway protects mice against the metabolic disturbances associated with diet-induced obesity (DIO) or leptin deficiency (*ob/ob*). Small molecule inhibition of the gut microbial enzyme choline TMA-lyase (CutC) does not reduce food intake, but is instead associated with beneficial remodeling of the gut microbiome, improvement in glucose tolerance, and enhanced energy expenditure. We also show that CutC inhibition is associated with reorganization of host circadian control of both phosphatidylcholine and energy metabolism. This study underscores the relationship between microbe and host metabolism, and provides evidence that gut microbe-derived trimethylamine (TMA) is a key regulator of the host circadian clock. This work also demonstrates that gut microbe-targeted enzyme inhibitors have untapped potential as anti-obesity therapeutics.

## Introduction

There is a growing body of evidence that microbes residing in the human intestine represent a critical environmental factor that influences virtually all aspects of human health and disease^5,6^. In fact, our gut microbiome plays a central role in vital processes such as energy harvest from our diet, entraining our immune system during early life, xenobiotic metabolism, and the production of a diverse array of small molecule metabolites that are essential to human life^5,6^. Although the microbiome field has uncovered many correlative relationships with human health and disease, there are few examples whereby alterations in the gut microbiome have been causally linked. One of the earliest and reproducible causal relationships established between gut microbes and human disease was is the link between alterations in gut microbial phyla (Bacteroidetes and Firmicutes) and obesity susceptibility, and the microbial transplantation studies revealing obesity susceptibility as a transmissible trait^1-4^. Specifically, obese mice harbor gut microbial communities with enhanced capacity to harvest energy from indigestible carbohydrates^3^, and transplantation of either cecal contents or feces from either obese mice or humans is sufficient to promote obesity and related insulin resistance in germ-free mouse recipients^1-4^. There is also accumulating evidence that antibiotic exposure in early life can predispose children to become overweight or obese later in life^7^, and antibiotic treatment in mice prior to weaning increases obesity and related insulin resistance in adulthood^8^. In fact, nearly all diseases where obesity is a predisposing comorbidity (diabetes, liver disease, cardiovascular disease, hypertension, chronic kidney disease, and diverse cancers) have been shown to have a clear gut microbiome link^9-14^.

Although there is now ample evidence that gut microbes play a contributory role in the development of obesity and related metabolic disorders, obesity-targeted drug discovery to this point has focused solely on targets encoded by the human genome. Our knowledge is rapidly expanding in regards to what types of microbes are associated with obesity and related disorders, including the repertoire of microbe-associated molecule patterns (MAMPs) they harbor and the vast array of metabolites that they produce. However, there are very few examples of where this information has been leveraged into clinically relevant therapeutics strategies. The microbiome-targeted therapeutic field has primarily focused on either pre-biotic or pro-biotic approaches, yet thus far these community restructuring approaches have resulted in very modest or non-significant effects in obesity-related disorders in human studies^15-19^. As an alternative microbiome-targeted approach, we and others have begun developing non-lethal selective small molecule inhibitors of bacterial enzymes with the hopes of reducing levels of disease-associated microbial metabolites^20-24^. In fact, we have recently shown that small molecule inhibition of the gut microbial transformation of choline into trimethylamine (TMA), the initial and rate-limiting step in TMAO generation^25^, can significantly reduce atherosclerosis, thrombosis, and adverse ventricular and kidney remodeling in mice^20-23^. In a day where host genetics/genomics approaches dominate, the discovery that the gut microbial TMAO pathway plays a contributory role in cardiovascular disease (CVD) risk highlights that the interplay between our diet and gut microbial enzymology play an extremely important role in modulating human disease susceptibility^25^. In fact, since 2011 the gut microbe-associated TMAO pathway has been associated with many human diseases associated with obesity including atherosclerosis^25,26^, thrombosis^27,28^, chronic kidney disease^29,30^, heart failure^31,32^, cancer^33,34^, and diabetes^35,36^. Given the fact that obesity is a comorbidity in all of these human diseases, here we set out to determine whether selective small molecule inhibition of the gut microbial choline transformation into TMA, a metabolic activity catalyzed by the microbial choline TMA lyase CutC^37^, can protect against metabolic disturbance in preclinical mouse models of obesity.

### Microbial choline TMA lyase inhibition protects mice from obesity development

To assess whether small molecule inhibition of gut microbial TMA production can protect mice from obesity, we treated mice with the non-lethal mechanism-based bacterial choline TMA lyase inhibitor iodomethylcholine (IMC)^20^. This small molecule inhibitor exhibits potent *in vivo* inhibition of the gut microbial choline TMA lyase enzyme CutC, lowering host plasma TMAO levels >90%^20,22,23^. Designed as a suicide substrate mechanism-based inhibitor, past studies reveal the vast majority of IMC is retained in bacteria and excreted in the feces with limited systemic exposure of the polar drug in the host^20^. When administered in a high fat diet (HFD), IMC effectively reduces levels of both the primary product of CutC TMA as well as the host liver-derived co-metabolite TMAO (Figure 1a,b). IMC was effective in blunting diet-induced obesity (DIO) (Figure 1c), without altering food intake (Fig. 1d). DIO mice treated with IMC also showed improvements in glucose tolerance (Figure 1e), and exhibited marked reductions in plasma insulin levels in fed mice but not under fasting conditions (Figure 1f). Next, we administered IMC to hyperphagic leptin-deficient (*ob/ob*) mice in a chow-based diet (Figure 1g-l). As expected, IMC effectively lowered both TMA and TMAO levels in *ob/ob* mice (Figure 1g,h). Quite unexpectedly, IMC dramatically protected *ob/ob* mice from body weight gain, where some of the drug treated mice were 15-20 grams lighter than control *ob/ob* mice (Figure 1i). Notably, IMC-driven protection from obesity was not associated with decreased food intake, which was paradoxically higher overall in IMC-treated mice (Figure 1j). Instead, IMC treatment promoted large increases in energy expenditure through both the light and dark cycles (Figure 1l, Figure 1 – figure supplement 1) Although IMC effectively reduced body weight and adiposity in *ob/ob* mice, this was not associated with improvements in glucose tolerance (Figure 1k). Given these striking results in DIO and *ob/ob* obesity progression models, we next wanted to determine if IMC could likewise provide obesity protection in mice with established obesity (i.e. a treatment regimen). To test this we fed C57BL/6J mice HFD for 6 weeks to allow for mice to reach obese conditions (body weights ranging from 35-40 grams) and then initiated IMC treatment. Although IMC did not promote weight loss, it did effectively reduce the trajectory of weight gain and improved glucose tolerance in this obesity treatment model (Figure 1 – figure supplement 1). During the early phases of this obesity treatment study, IMC-treated mice again exhibited unexpected increased food intake compared to controls (Figure 1 – figure supplement 1). To more comprehensively understand the global effects of choline TMA lyase inhibition on white adipose tissue gene expression, we performed unbiased RNA sequencing (Figure 1 – figure supplement 1). Principal coordinate analysis (PCA) of RNA expression profiles showed clear separation of adipose gene expression between HFD control versus HFD with IMC (Figure 1 – figure supplement 1). The most differentially expressed genes altered by IMC were enriched in common pathways of adipogenesis, fatty acid metabolism, inflammation, and cytokine signaling (Figure 1 – figure supplement 1). Collectively, these data demonstrate that gut microbe-targeted choline TMA lyase inhibition can protect mice from obesity and selectively reorganize host adipose tissue gene expression.

**Figure 1.**
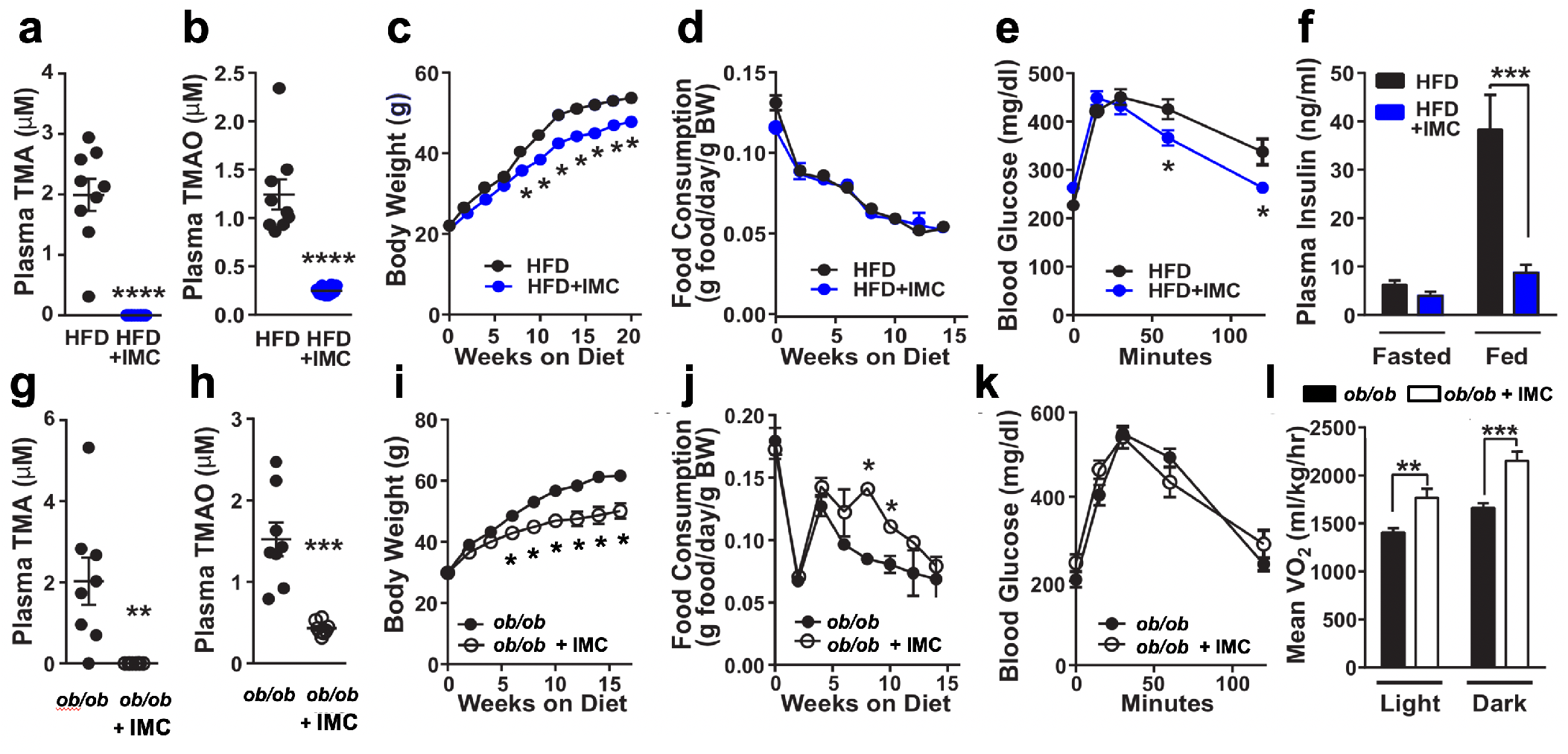
Small Molecule Choline TMA Lyase Inhibition Improves Obesity. Panels **a**-**f** and **g**-**l** represent data from control and IMC-treated HFD-fed and *ob/ob* mice, respectively. **a** and **g**, plasma TMA levels. **b** and **h**, plasma TMAO levels. **c** and **i**, biweekly body weights. **d** and **j**, biweekly food consumption. **e** and **k**, glucose tolerance test. **f**, plasma insulin levels. **l**, average oxygen consumption. In panels **a**-**e** and **g**-**k**, groups were compared using t-tests. In panels **f** and **l**, groups were compared using two-way ANOVA with Tukey’s multiple comparisons test. Significance is defined as: *, P<0.05. **, P<0.01. ***, P<0.001, ****, P<0.0001. n = 6-10 per group.

### Choline TMA lyase inhibition promotes beneficial remodeling of the gut microbiome

One theoretical advantage of non-lethal microbe-targeted choline TMA lyase inhibitors, compared to antibiotic therapies, is that microbiome restructuring effects of the drug are expected to be primarily target the species that rely on choline as a carbon or nitrogen source. Therefore, we next examined whether IMC treatment was associated with alterations in choline utilizers and other members of the gut microbiome community that may contribute to improvement in obesity-related metabolic disturbances. IMC treatment in HFD-fed mice resulted in significant restructuring of the cecal microbiome at every taxonomic level (Figure 2a-e). PCA of microbial taxa revealed distinct clusters, indicating that IMC promoted restructuring of the cecal microbiome (Figure 2a). At the phylum level, HFD-fed IMC-treated mice had large increases in *Verrucomicrobia* and *Bacteroidetes*, and significant reductions in *Firmicutes* (Figure 2b,c). It is important to note that the ratio of *Firmicutes* to *Bacteroidetes*, which has been repeatedly associated with obesity in both humans and mice^1-4^, is significantly reduced by IMC treatment of the HFD-fed mice (Figure 2c). Performance of linear discriminant analysis coupled with effect size measurements (LEfSe analysis) revealed that IMC promoted significant reductions in the proportions of the taxa *Desulfovibrionaceae, Butyrovibrio, Bilophia, Peptococcaceae*, and *Dorea*, and increases in *Odoribacter, Roseburia*, and several members of the *Clostridiaceae* family (Figure 2d). We also examined whether the proportions of cecal genera were significantly correlated with plasma TMA, body weight, and glucose tolerance across in DIO mice, and found that increases in *Akkermansia* and reductions in *Bilophila* were significantly correlated with plasma TMA level, and similar yet non-significant trends were found with body weight and glucose tolerance (Figure 2e). Similar to HFD-fed mice (Figure 2a-e), IMC also significantly altered the cecal microbiome in *ob/ob* mice (Figure 2f-2i). In *ob/ob* mice, IMC significantly reduced the proportions of *Adlercreutzia, Rikenellaceae, Peptococcaceae*, and *Clostridiales*, while increasing levels of *Molicutes, Allobaculum, Peptostreptococcaceae*, and *Akkermansia* (Figure 2g-h). Correlation analysis in *ob/ob* mice revealed that IMC-induced alterations in *Akkermansia, Clostridiales*, and *Allobaculum* were significantly associated with circulating TMA and body weight (Figure 2i). Notably, in both the *ob/ob* and DIO models, the large increase in the proportions of *Verrucomicrobia* phylum seen with IMC are largely explained by increased level of *Akkermansia mucinophila*, which has been pursued as a therapeutic probiotic strain for diabetes amelioration in other studies^38-40^, and is correlated to TMA and body weight phenotypes in the current study. Collectively, these data demonstrate that inhibition of gut microbial choline to TMA transformation with a selective non-lethal small molecule inhibitor promotes beneficial restructuring of the gut microbiome that may contribute in part to improvements in energy metabolism and obesity observed in the host.

**Figure 2.**
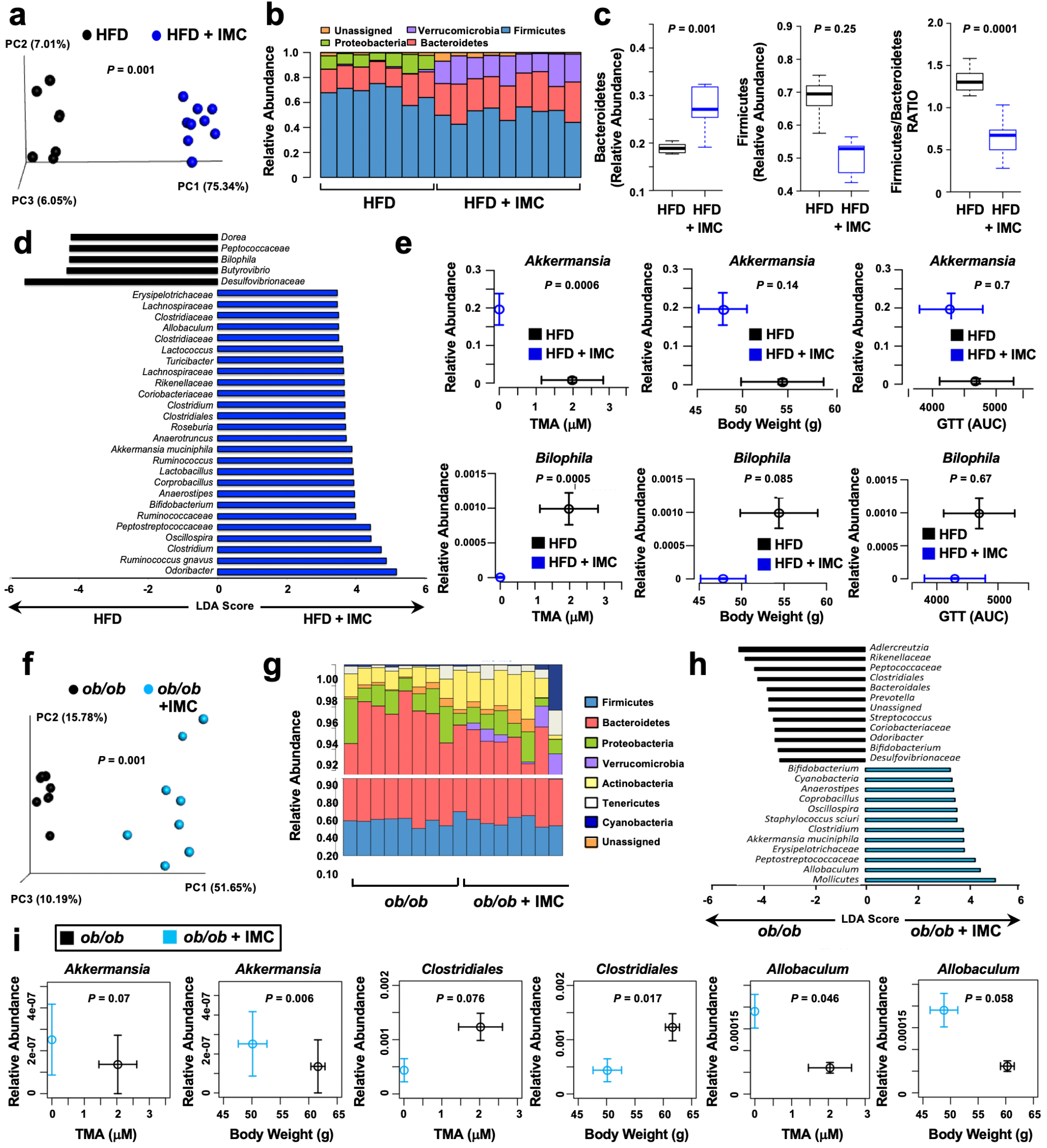
Small Molecule TMA Lyase Inhibition Promotes Beneficial Remodeling of the Gut Microbiome. Panels **a**-**e** and **f**-**i** represent data from control and IMC-treated HFD-fed and *ob/ob* mice, respectively. **a** and **f**, Principal co-ordinate analysis plot of microbiota profiles built from weighted Unifrac distances. Each point represents a single sample from a single mouse. Positions of points in space display dissimilarities in the microbiota, with points further from one another being more dissimilar. **b** and **g**, Barplot of cecal microbiota at the phylum level. Each bar represents an individual mouse and each color an individual phyla. **c**. Relative abundance of Firmicutes, Bacteroidetes and the Firmicutes to Baceteroidetes ratio. The boxes represent the 25^th^ and 75^th^ quartiles, and the line displays the median value within each group. Points extending beyond the lines are outliers defined as values greater or less than 1.5 times the interquartile range. **d** and **h**, Linear discriminatory analyses plot of taxa differing significantly with IMC treatment. **e and i**, Correlation between taxa and plasma TMA levels, body weight, and glucose tolerance (HFD-fed mice only). Values in both X and Y directions are plotted as mean ± SEM.

### The gut microbial TMAO pathway regulates host circadian rhythms

Although there is clear evidence that TMAO can directly impact cell signaling in macrophages^41^, endothelial cells^41^, and platelets^27^, it is still incompletely understood how the TMAO pathway is mechanistically linked to all of these common obesity-related diseases. Because of the recent appreciation of the gut microbiome as a coordinator of host metabolism and obesity-related phenotypes^40-43^, we hypothesized that the metaorganismal TMAO pathway regulates host circadian rhythms to impact obesity and its associated cardiometabolic complications. The data that led us to this possibility includes: (1) misalignment of the host circadian clock is associated with the same human diseases that the TMAO pathway has been linked to^40-43^, (2) the gut microbiome oscillates in a circadian manner and this is highly gender specific (similar to the TMAO pathway)^40-42^, and (3) disruption of the host circadian core clock machinery promotes gut microbial “dysbiosis” and alters circulating levels of gut-derived metabolites^40-43^. Given these clear overlaps, we investigated whether different substrates and regulatory genes within the TMAO pathway exhibited circadian rhythmicity. Indeed, we found that several nodes within the gut microbial TMAO pathway showed strong diurnal oscillations in mice (Figure 3a). Plasma levels of both choline, carnitine, and TMA were relatively low during the light cycle, and peak at the beginning of the dark cycle when mice are most active and consuming food (Figure 3a). However, TMAO showed more modest oscillatory behavior (Figure 3a and Figure 3 – figure supplement 1). We also found that the hepatic expression of flavin-containing monooxgygenase 3 (*Fmo3*) is quite low during the day, but increases 6-fold during the early dark cycle (Figure 3a). However, this pattern for *Fmo3* expression is opposite in white adipose tissue (Figure 4 – figure supplement 1). Unexpectedly, we identified a tissue-specific circadian rhythm in the host TMA receptor (trace amine-associated receptor 5; *Taar5*) (Figure 3a). It is important to note that *Taar5* oscillation specifically occurs in skeletal (Figure 3a; Figure 3 – figure supplement; Figure 4 – figure supplement 1) and cardiac muscle (Figure 4 – figure supplement 1), and shows no oscillatory behavior in the olfactory bulb (Figure 4 – figure supplement 1), where it is important for sensing the “fishy odor” smell of TMA^44,45^. Collectively, these data suggest that the gut microbial TMAO pathway dynamically oscillates in a circadian manner.

**Figure 3.**
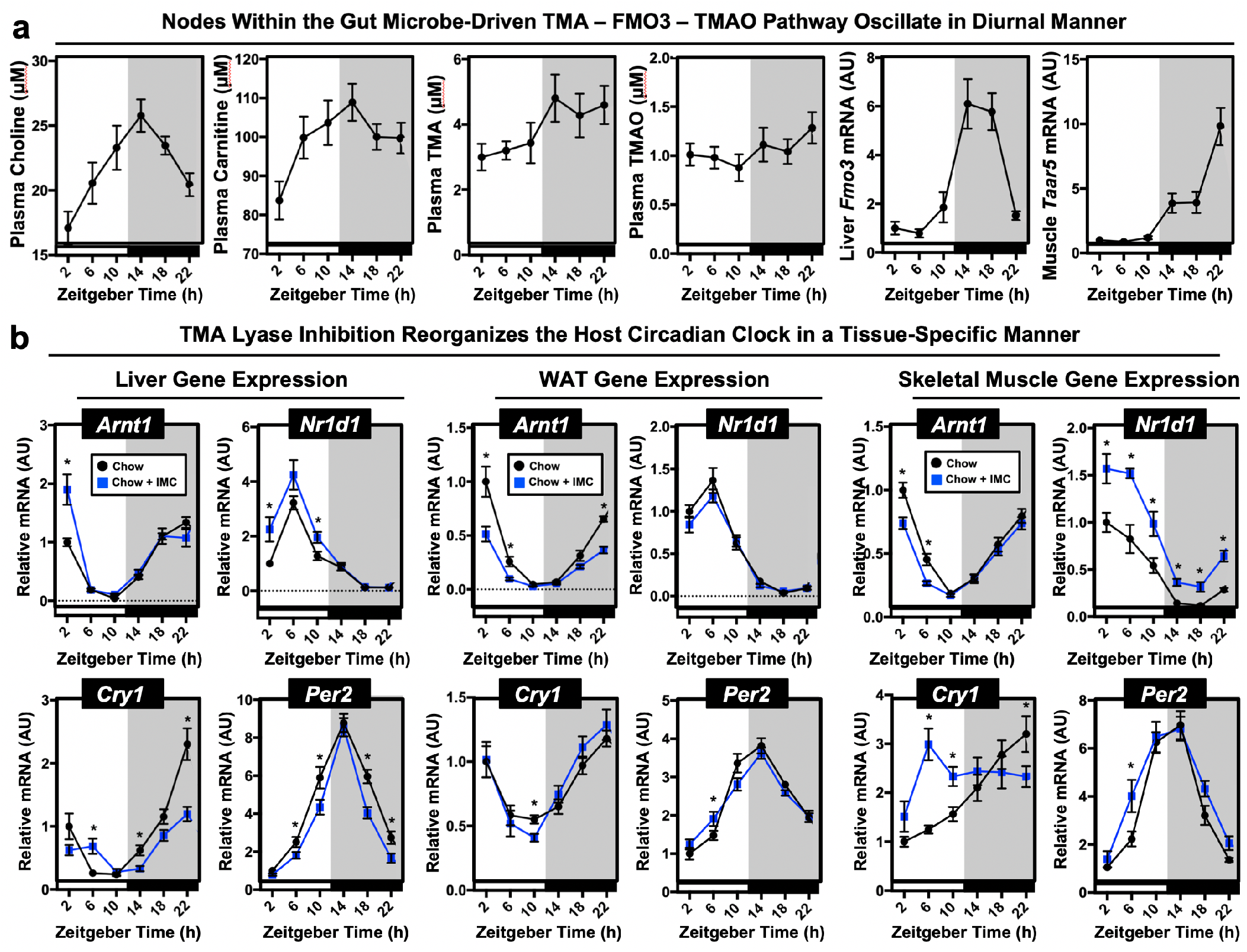
Nodes Within the Metaorganismal TMAO Pathway Show Diurnal Rhythmicity and Choline TMA Lyase Inhibition Alters the Host Circadian Clock. Wild type male C57BL/6J mice were necropsied at 4 hour intervals to collect plasma and tissue over a 24 hour period. **a**. Plasma choline, L-carnitine, trimethylamine (TMA); trimethylamine N-oxide (TMAO) were quantified by LC-MS/MS; hepatic expression of *Fmo3* mRNA or gastrocnemius muscle expression of *Taar5* mRNA was quantified via qPCR (n = 7-8). **b**. Wild type male C57BL/6J mice were fed chow or chow supplemented with the choline TMA lyase inhibitor iodomethylcholine (IMC) for 7 days. Mice were then necropsied at 4 hour intervals to collect liver, gonadal white adipose tissue, or gastrocnemius skeletal muscle. We then performed qPCR to examine the mRNA expression in liver, white adipose, and gastrocnemius skeletal muscle of key circadian clock regulators including aryl hydrocarbon receptor nuclear translocator like (*Arntl;* BMAL1), nuclear receptor subfamily 1 group D member 1 (*Nr1d1*; RevErb*α*), cryptochrome 1 (*Cry1*), or period 2 (*Per2*) (n = 7-8). Significance (*p* <0.05) between diet groups at specified Zeitgeber (ZT) time points were compared using Student’s t-tests.

**Figure 4.**
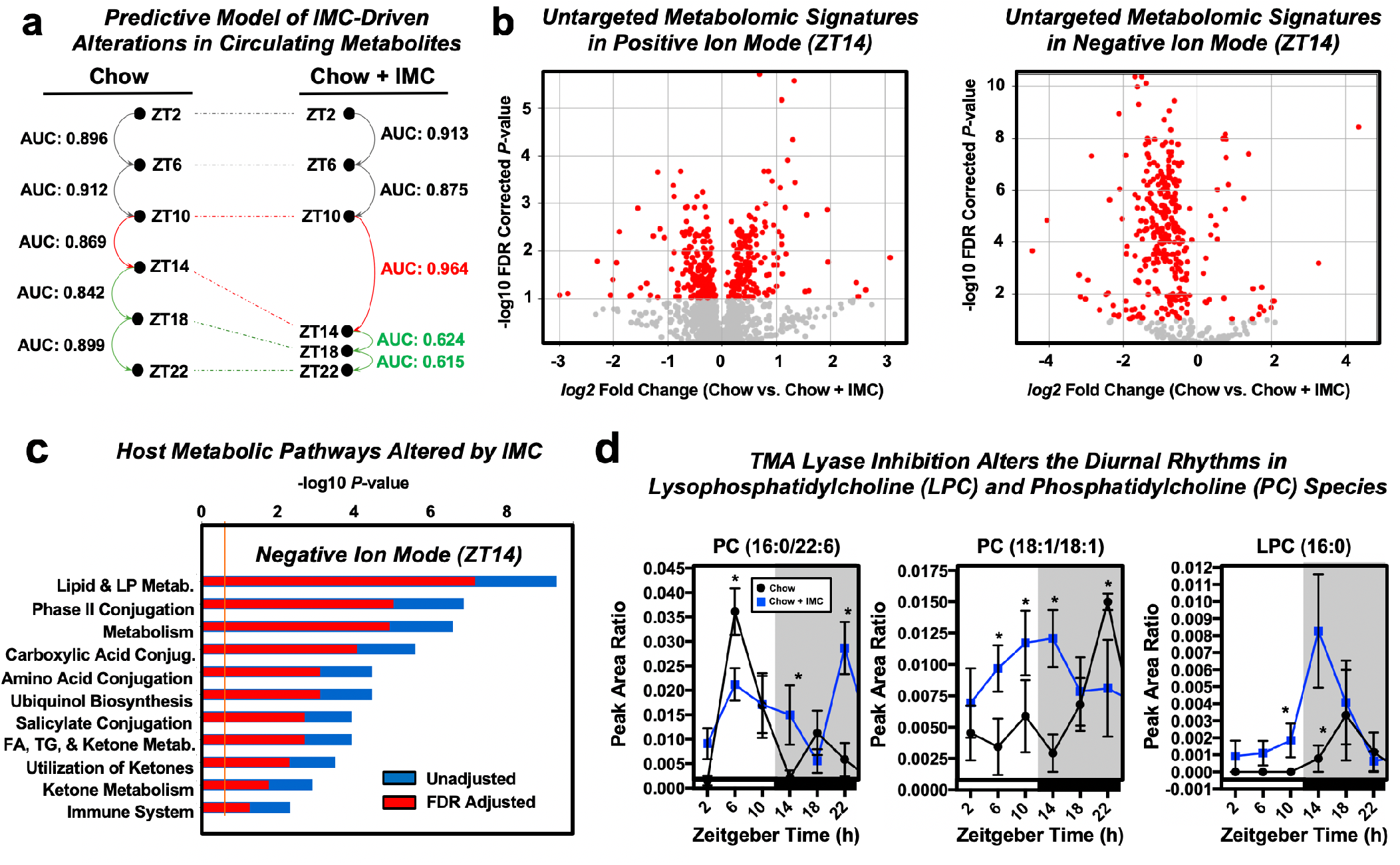
Small Molecule Inhibition of Choline TMA Lyases Impacts Circadian Oscillations in Choline Containing Phospholipids in the Host Circulation. Wild type male C57BL/6J mice were fed chow or chow supplemented with the choline TMA lyase inhibitor iodomethylcholine (IMC) for 7 days. Mice were then necropsied at 4 hour intervals to collect blood and tissue, and plasma time points were subjected to LC-MS/MS-based untargeted metabolomics (n = 7-8). **a**. Selective Paired Ion Contrast Analysis (SPICA)^51^ was used to identify time points where the plasma metabolome was most dramatically altered by IMC treatment compared to zeitgeber-matched controls. Pairwise analysis was conducted between all adjacent timepoints for the control and IMC treatment groups, resulting in a total of 10 comparisons made (5 for Chow and 5 for Chow + IMC). Differences in the global plasma metabolome for each pairwise comparison were quantified via receiver operating characteristic (ROC) curve construction and area under the curve (AUC) calculations via Monte Carlo cross validation procedures in SPICA^51^ revealed that IMC most dramatically altered the plasma metabolome during the transition from ZT10 to ZT14. As a result, subsequent data analyses focused on this T10 to T14 transition, which also coincided with the light to dark phase transition and when the mice began to eat. **b**. Volcano plots of the significantly altered metabolites (red) in positive and negative ion mode at the ZT14 time point. **c**. Pathway analysis of data collected in negative ion mode at ZT14 reveals that IMC alters host lipid and lipoprotein metabolism among other metabolic pathways. **d**. Relative plasma levels of phosphatidylcholine (PC) and lysophosphatidylcholine (LPC) species are altered by IMC.

### Gut microbe-targeted inhibition of TMA production rewires host-microbe metabolic crosstalk

To determine whether TMA or TMAO could potentially serve as a gut microbe-derived signal to entrain the host circadian clock we treated a set of mice with IMC acutely (7 days) and examined effects on the core clock machinery as well as clock-mediated regulation of host metabolism. In this circadian study, IMC treatment effectively lowered circulating TMA and TMAO at every time point, yet there was still a highly rhythmic low level of plasma TMA and TMAO observed, possibly originating from microbial metabolism of non-choline (i.e. carnitine, *γ*-butyrobetaine, betaine, trimethyllysine) nutrient sources via microbial enzymes that are not inhibited by IMC^46-48^ (Figure 3 – figure supplement 1). Unexpectedly, Provision of IMC altered the expression of core clock transcription factors in a highly tissue-specific manner. In the liver, microbial choline TMA lyase inhibition modestly increased the peak amplitude of the core clock transcription factors *Bmal1* and *Rev-erbα*, yet blunted the rhythmic expression of *Cry1* (Figure 3b). In white adipose tissue, choline TMA lyase inhibition blunted the peak amplitude of *Bmal1* (Figure 3b). However, in skeletal muscle we found the most striking differences in the core clock machinery where IMC-treated mice showed marked increases at all time points for *Rev-erbα* and a near complete phase shift for *Cry1* (Figure 3b; Figure 3 – figure supplement 1). Prior studies have confirmed that IMC is poorly absorbed in the host and selectively accumulates in gut microbiota^20,22^. It is tempting to speculate that the circadian oscillations in skeletal muscle are so drug responsive due in part to the fact that the TMA receptor *Taar5* oscillates there (Figure 3a; Figure 3 – figure supplement 1) to sense TMA and is not a “direct” result of IMC *per se*, but rather IMC effect on gut microbiota production of TMA. It is important to note that IMC does not alter the expression of core clock genes in area of the central nervous system like the olfactory bulb where *Taar5* is abundantly expressed throughout the day and does not oscillate diurnally.

Recent literature has demonstrated that the composition of the gut microbiome is also under circadian regulation^40-42^. These reports show that even at the phylum level, bacterial communities oscillate in a highly circadian manner, and gut microbe community oscillation can impact host metabolic rhythms that are linked to human disease^40-42^. However, there is almost nothing known regarding what regulates this diurnal rhythm in gut microbial communities. Here we demonstrate that microbial production of TMA, as modulated by the choline TMA lyase inhibitor IMC, has striking effects on circadian oscillations in gut microbial communities (Figure 2 – figure supplement 1). PCA analyses revealed distinct group clusters, indicating that IMC promoted restructuring of the cecal microbial community composition in a circadian manner (Figure 2 – figure supplement 1). At the phylum level, IMC treatment results in large relative increases in *Bacteroidetes*, while lowering the overall levels of *Firmicutes* at all time points (Figure 2 – figure supplement 1). Interestingly, IMC treatment promoted a significant increase in the abundance of *Akkermansia mucinophila*, proportions of which have been reported to be inversely correlated with body weight and insulin sensitivity in numerous studies in both rodents and humans^38-40^. In addition, IMC treatment increased the level of S24-7 Bacteroidetes family members, which have recently been reported to be depleted in several mouse models of obesity-related metabolic disturbance^49-51^. Collectively, these data demonstrate that gut microbial production of TMA may be an important cue to impact the diurnal oscillations in gut microbial community structure and metabolic function in the host.

In an attempt to understand how choline TMA lyase inhibition altered host metabolic processes in an unbiased way, we performed untargeted plasma metabolomics across a 24-hour period in IMC-treated mice (Figure 4). This large circadian metabolomics dataset was initially evaluated for any large-scale trends utilizing Selective Paired Ion Contrast Analysis (SPICA)^52^ to identify key zeitgeber (ZT) time points where IMC elicited the most significant alterations in the plasma metabolome (Figure 4a). Although there were clear IMC-induced metabolomic alterations at every time point, the most significant alterations were seen during the ZT10 to ZT14 transition, which coincides with the light to dark transition when mice typically begin to forage for food. Focusing on the ZT14 time point, we observed a large number of differentially abundant metabolites in the IMC-treated group that were in common pathways of lipid and lipoprotein metabolism, phase II conjugation, and other various macronutrient metabolic pathways (Figure 4b,c). In particular, we found that choline TMA lyase inhibition was associated with alterations in diurnal rhythms of phosphatidylcholine (PC) co-metabolites (Figure 4d). In fact, gut microbial TMA lyase inhibition caused near complete circadian phase shifts for two PC species (16:0/22:6 and 18:1/18:1), and two saturated species (16:0 and 18:0) of lysophosphatidylcholine (LPC), which showed robust oscillatory behavior distinct from control mice (Figure 4d). Furthermore, we found that the oscillatory behavior of genes involved in PC synthesis including choline kinase α (CKα) and phosphatidylethanolamine methyltransferase (PEMT) were significantly altered in TMA lyase inhibited mice (Figure 3 – figure supplement 1). It is important to note that IMC did not alter key plasma hormones (corticosterone or insulin) or lipids (non-esterified fatty acids) that are known to have strong effects on phospholipid metabolism themselves (Figure 4 – figure supplement 1). Collectively, these data suggest that gut microbial TMA production from the choline TMA lyase CutC is a critical determinant of host circadian rhythms in PC homeostasis.

## Discussion

Although obesity drug discovery has historically targeted pathways in the human host, a fertile period in biomedical research lies ahead where we instead we target the microorganisms that live within us to improve obesity and related disorders. This paradigm shift in drug discovery is needed in light of the clear and reproducible associations between the gut microbiome and almost every human disease. Now we are faced with both the challenge and opportunity to test whether microbe-targeted therapeutic strategies can improve health in the human metaorganism without negatively impacting the symbiotic relationships that have co-evolved. Although traditional microbiome manipulating approaches such as antibiotics, prebiotics, probiotics, and fecal microbial transplantation have shown their own unique strengths and weaknesses, each of these present unique challenges particularly for use in chronic diseases such as obesity. As we move toward selective nonlethal small molecule therapeutics, the hope is to have exquisite target selectivity and limited systemic drug exposure given the targets are microbial in nature. This natural progression parallels the paradigm shifts in oncology which have transitioned from broadly cytotoxic chemotherapies to target-selective small molecule and biologics-based therapeutics. Here we provide the first evidence that a selective nonlethal inhibitor to the microbial choline TMA lyase CutC can have beneficial effects on host energy metabolism, adiposity, and insulin sensitivity. The current study demonstrates that gut microbe-targeted suppression of choline to TMA generation: (1) protects the host from diet-induced obesity and glucose intolerance, (2) protects *ob/ob* mice from obesity, (3) increases energy expenditure and alters the expression of lipid metabolic genes in white adipose tissue, and (4) beneficially reorganizes gut microbial communities to improve the obesity related *Firmicutes* to *Bacteroidetes* ratio and increase abundance of *Akkermansia mucinophila*. This study also reveals that nodes within the gut microbial TMAO pathway exhibit circadian oscillatory behavior, and that inhibition of microbial choline to TMA transformation is associated with alterations in: (1) the expression of core clock machinery in metabolic tissues; (2) the diurnal rhythmic behavior of gut microbial communities; and (3) circadian oscillation in choline-containing phospholipids. Collectively, our findings reinforce the notion that the gut microbial TMAO pathway is a strong candidate for therapeutic intervention across a spectrum of obesity-related diseases, and have uncovered metaorganismal crosstalk between gut microbe-derived TMA and circadian metabolic oscillations in mice. Interestingly, during the preparation of this manuscript a recent meta-analysis of clinical studies demonstrated that circulating TMAO levels are dose-dependently associated with obesity in humans^53^, highlighting the potential translation of this work.

As drug discovery advances in the area of small molecule nonlethal bacterial enzyme inhibitors it is key to understand how these drugs impact microbial ecology in the gut and other microenvironments. As we have previously reported^20,22,23^, IMC treatment does promote a significant remodeling of the cecal microbiome in mice. These IMC-induced alterations occur as early as one week (Figure 2 – figure supplement 1) and persist even after twenty weeks (Figure 2). It is important to note that the observed IMC-induced alterations in the gut microbiome are generally expected to be beneficial. For instance, IMC lowers the ratio of Firmicutes to Bacteroidetes (Figure 2c), which is known to be elevated in obese humans and rodents^1-4^. Of the taxa that were significantly correlated with circulating TMA levels (*Akkermansia, Bilophila, Clostridiales*, and *Allobaculum*), only *Akkermansia* was likewise correlated with body weight in *ob/ob* mice (Figure 2i). It is important to note that previous studies have shown beneficial effects of *Akkermansia mucinophila* in obesity and diabetes, which has prompted its development as a probiotic^36-38^. In fact supplementation with *Akkermansia mucinophila* can improve glucose tolerance in obese mice^38-40^. It is important to note that increases in *Akkermansia mucinophila* has been consistently seen in mice treated with TMA lyase inhibitor^20,22,23^, suggesting that small molecule inhibition of gut microbial choline TMA lyase inhibitors may be an alternative means to enrich gut microbiomes with *Akkermansia mucinophila* in addition to developing probiotic approaches. As small molecule bacterial enzymes inhibitors are developed it will be extremely important to understand their effects on microbial ecology, and it is expected that some of the beneficial effects of these drugs will indeed originate from the restructuring of gut microbiome communities. In fact, this is not an uncommon mechanism by which host targeted drugs impact human health. A recent study showed that nearly a quarter of commonly used host-targeted drugs have microbiome-altering properties^54^, and in the context of diabetes therapeutics it is important to note metformin’s anti-diabetic effects are partially mediated by the drug’s microbiome altering properties^55^. Given the strong association between gut microbiome and obesity and diabetes^1-4^, it will likely be advantageous to find therapeutics that beneficially remodel the gut microbiome as well as engage either their microbe or host target of interest. One burgeoning area of microbe-host cross talk relevant to human disease is the intersection between gut microbes and host circadian rhythms^40-43^. It is well appreciated that a transcriptional-translational feedback loop (TTFL) exists in mammalian cells to orchestrate an approximately 24-hour oscillatory rhythm in the expression of thousands of genes^40-43^. The mammalian clock is coordinated by core transcription factors CLOCK and BMAL1, which peak during light phases, and crytochromes (CRYs) and period genes (PERs), which are most active during dark phases. Under normal conditions, the clock regulated TTFL maintains cell autonomous homeostatic responses to environmental “zeitgebers” including light, food, xenobiotics, and exercise^40-43^. However, disruption of normal circadian rhythms induced by abnormal light exposure, sleep-activity, or eating patterns has been associated with the development of many human diseases including obesity, diabetes, CVD, kidney disease, cancer, and neurological disease^40-43^. Therefore, “chronotherapies” or therapies that prevent circadian disruption hold promise across a wide spectrum of diseases. Interestingly, it has been discovered that circadian disruption is associated with a marked reorganization of gut microbial communities, and microbial abundance in the gut exhibits circadian rhythmicity^40-43^. However, whether gut microbial metabolites contribute to circadian disruption is not well understood. Here we show that multiple nodes with the metaorganismal TMAO pathway (choline, TMA, *Fmo3*, and *Taar5*) oscillate in a circadian manner, and that inhibition of the choline TMA lyase CutC/D rewires the host circadian clock itself, as well as circadian rhythms in the cecal microbiome, as well as clock-driven reorganization of host lipid metabolic processes. The impact of gut microbial choline TMA lyase inhibition on the expression of core clock genes is modes in metabolic tissues such as the liver or white adipose tissue (Figure 3b; Figure 3 – figure supplement 1). However, in skeletal muscle, where the host TMA receptor *Taar5* exhibits oscillatory behavior (Figure 3a), IMC treatment elicits profound alterations in the expression of *Nr1d1* (RevErb*α*) and cryptochrome 1 (*Cry1*). The oscillatory behavior of the TMA receptor *Taar5* is specific to both skeletal and cardiac muscle, and is not seen in the olfactory bulb where *Taar5* is essential to sense the fish-like odor of TMA^44,45^. A recent report demonstrated that TMAO, but not TMA, can bind to and activate the endoplasmic reticulum stress (ER stress)-related kinase PERK (*EIF2AK3*), and that TMAO binding to PERK regulates the expression of the forkhead transcription factor *FoxO1*^56^. It is interesting to note that PERK was recently shown to regulate circadian rhythms that support sleep/wake patterns and cancer growth^57,58^. We also observed tissue-specific alterations in the circadian expression of *FoxO1* with IMC treatment (Figure 3 – figure supplement 1; Figure 4 – figure supplement 1). However, additional studies are needed to understand whether the TMAO-PERK-FOXO1 signaling axis is mechanistically linked to oscillatory behavior in metabolism or associated metabolic disease. It is interesting to note that several independent metabolomics studies have found that circulating levels of TMAO exhibit circadian oscillatory behavior^59-61^. In a similar manner, the expression of flavin-containing monooxygenase enzymes which convert TMA to TMAO are also under direct circadian clock transcriptional regulation^62-64^. Collectively, these emerging data suggest that gut microbe-derived TMA and potentially its metabolic co-metabolite TMAO may be underappreciated microbe-derived signals that impact host circadian rhythms in metabolism. Further studies are now warranted to test whether TMA lyase inhibitors of the host liver TMAO-producing enzyme FMO3 can act as chronotherapies to improve circadian misalignment-associated diseases.

The metaorganismal TMAO pathway represents only one of many microbial metabolic circuits that have been associated with human disease. In fact, many microbe-associated metabolites such as short chain fatty acids, secondary bile acids, phenolic acids, polyamines, among others have been associated with many human diseases^5,11^. In a circadian and/or meal-related manner gut microbes produce a diverse array of metabolites that reach micromolar to millimolar concentrations in the blood, making the collective gut microbiome an active endocrine organ^5^. Small molecule metabolites are well known to be mediators of signaling interactions in the host, and this work provides evidence that diet-microbe-host metabolic interplay that can shape normal diurnal rhythms in metabolism and obesity susceptibility in mice. Our work, that of many others, demonstrates that there is clear evidence of bi-directional crosstalk between the gut microbial endocrine organ and host metabolism. As drug discovery advances it will be important to move beyond targets based solely in the human host. This work highlights that non-lethal gut microbe-targeted enzyme inhibitors can serve as effective anti-obesity therapeutics in preclinical animal models, and provides proof of concept that this may be a generalizable approach to target metaorganismal crosstalk in other disease contexts. In fact, selective mechanism-based inhibition of bacterial enzymes has the advantage over host targeting given that small molecules can be designed to avoid systemic absorption and exposure, thereby minimizing potential host off target effects. As shown here with the gut microbial TMAO pathway, it is easy to envision that other microbe-host interactions are mechanistically linked to host disease pathogenesis, serving as the basis for the rational design of microbe-targeted therapeutics that improve human health.

## METHODS

### Mice and Experimental Diets

C57Bl6/J and *ob/ob* mice were purchased from The Jackson Laboratory (Bar Harbor, ME) and treated with a gut microbe-targeted small molecule inhibitor of choline TMA lyase called iodomethylcholine (IMC)^20^. For high fat diet feeding experiments, C57Bl6/J were maintained on 60% high fat diet alone or with the addition of 0.06%w/w IMC (D12942 and custom diet D15102401, respectively, Research Diets, Inc.). For *ob/ob* and circadian studies, mice were maintained on a standardized control chow diet (TD:130104, Teklad) with or without supplemental with IMC (0.06% w/w). Body weight was measured weekly. Food intake was assessed by weighing the food consumed weekly divided by the number of mice in each cage and normalized to the average body weight. For circadian rhythm studies, 9-week old C57Bl6/J male mice were adapted for 2 weeks on a standardized minimal choline chow (Envigo diet # TD.130104) with a strict 12 h:12 h light:dark cycle after shipment. Mice were then maintained on standardized chow (Envigo diet # TD.130104) or the same diet supplemented with 0.06% IMC (Envigo diet # 150813). After 7 days, plasma and tissue collection were performed every 4 hours over a 24-hour period. All dark cycle necropsies were performed under red light conditions. In the obesity treatment study paradigm C57Bl6/J mice fed a high fat diet (Research Diets # D12942) for 6 weeks to establish obesity (body weight > 35 grams), and after 6 weeks of diet-induced obesity mice were continued on HFD alone (Research Diets # D12942) or the same HFD containing IMC for another 10 weeks to test whether IMC can improve obesity-related phenotypes. For all studies plasma was collected by cardiac puncture, and liver, WAT, and skeletal muscle was collected, flash frozen, and stored at −80°C until the time of analysis. All mice were maintained in an Association for the Assessment and Accreditation of Laboratory Animal Care, International-approved animal facility, and all experimental protocols were approved by the Institutional Animal Care and use Committee of the Cleveland Clinic.

### Synthesis of Iodomethylcholine (IMC) Iodide

Iodomethylcholine iodide was prepared using a previously-reported method using 2-dimethylethanolamine and diiodomethane as reactants in acetonitrile followed by recrystallization from dry ethanol^66. 1^H- and ^13^C-NMRs of IMC were both consistent with that in the reported literature^66^, as well as consistent based on proton and carbon chemical shift assignments indicated below. High resolution MS corroborated the expected cation mass and provided further evidence of structural identity.

^1^H-NMR (600 MHz, D_2_O): *δ* 5.13 (s, 2H, -N-C<u>H</u>_2_-I), 3.90 (t, J = 4.8 Hz, 2H, -CH_2_-C<u>H</u>_2_-OH), 3.52 (t, J = 4.8 Hz, 2H, -N-C<u>H</u>_2_-CH_2_-), 3.16 (s, 6H, -N(C<u>H</u>_3_)_2_);

^13^C-NMR (150 MHz, D_2_O): *δ* 65.7 (-CH_2_-<u>C</u>H_2_-OH), 55.4 (-N-<u>C</u>H_2_-CH_2_-), 52.4 (-N(<u>C</u>H_3_)_2_), 32.3 (-N-<u>C</u>H_2_-I);

HRMS (ESI/TOF): m/z (M^+^) calculated for C_5_H_13_INO, 230.0036; found, 230.0033.

### Measurement of Plasma Trimethylamine (TMA) and Trimethylamine-N-oxide (TMAO)

Stable isotope dilution high performance liquid chromatography with on-line tandem mass spectrometry (LC–MS/MS) was used for quantification of levels of TMAO, TMA, choline, carnitine, and *γ-butyrobetaine* in plasma, as previously described^67^. Their d9(methyl)- isotopologues were used as internal standards. LC–MS/MS analyses were performed on a Shimadzu 8050 triple quadrupole mass spectrometer. IMC and d2-IMC, along with other metabolites, were monitored using multiple reaction monitoring of precursor and characteristic product ions as follows: m/z 230.0 → 58.0 for IMC; m/z 232.0 → 60.1 for d2-IMC; m/z 76.0 → 58.1 for TMAO; m/z 85.0 → 66.2 for d9-TMAO; m/z 60.2 → 44.2 for TMA; m/z 69.0 → 49.1 for d9-TMA; m/z 104.0 → 60.1 for choline; m/z 113.1 → 69.2 for d9-choline; m/z 118.0 → 58.1 for betaine; m/z 127.0 → 66.2 for d9-betaine.

### Analysis of Gene Expression in Mouse Tissues

RNA was isolated via the RNAeasy lipid tissue mini kit (Qiagen) from multiple tissues. RNA samples were checked for quality and quantity using the Bio-analyzer (Agilent). RNA-SEQ libraries were generated using the Illumina mRNA TruSEQ Directional library kit and sequenced using an Illumina HiSEQ4000 (both according to the Manufacturer’s instructions). RNA sequencing was performed by the University of Chicago Genomics Facility. Raw sequence files will be deposited in the Sequence Read Archive before publication (SRA). Paired-ended 1050 bp reads were trimmed with Trim Galore (v.0.3.3, http://www.bioinformatics.babraham.ac.uk/projects/trim_galore) and controlled for quality with FastQC (v0.11.3, http://www.bioinformatics.bbsrc.ac.uk/projects/fastqc) before alignment to the Mus musculus genome (Mm10 using UCSC transcript annotations downloaded July 2016). Reads were aligned using the STAR alignerSTAR in single-pass mode (v.2.5.2a_modified, https://github.com/alexdobin/STAR)^68^ with standard parameters but specifying ‘–alignIntronMax 100000 –quantMode GeneCounts’. Overall alignment ranged from 88–99% with 61–75% mapping uniquely. Transcripts with fewer than one mapped read per million (MMR) in all samples were filtered out before differential expression (DE) analysis. The filtering step removed 12,692/24,411 transcripts (52%). Raw counts were loaded into R (http://www.R-project.org/) (R Development Core Team, 2015) and edgeR^69^ was used to perform upper quantile, between-lane normalization, and DE analysis. Values generated with the cpm function of edgeR, including library size normalization and log2 conversion, were used in figures. Heat maps were generated of top 50 differentially expressed transcripts using pheatmap^70^. Reactome-based pathway analysis was performed using an open-sourced R package: ReactomePA^71^. For real time quantitative PCR (qPCR) analyses ∼20 mg of snap-frozen liver tissue was homogenized in the 1-mL TRIzol reagent (Thermo Fisher Scientific, Cat. No. 15596018). Furthermore, 200 mL of chloroform were added and samples were spun down at 13,000 revolutions/min for 5 min. Upper clear layer were passed through the RNeasy Mini Spin Columns (Cat. No. 74104) for clean up of RNA according to manufacturer’s instructions. DNase treatment (10 U/reaction) was performed according to the Qiagen RNeasy kit (Cat. No. 74104). Concentrations of high purity RNA were measured using Nanodrop (Thermo Fisher Scientific, ND-2000). Reverse transcription to generate cDNA was performed using qscript mastermix (Quanta-Bio Cat. No. 101414-106) as recommended by the manufacturer using 750 ng of RNA template. Resulting cDNA was diluted 10x and used in the real-time PCR reaction using an Applied Biosystems Step One Plus thermocycler. Relative mRNA levels were calculated based on the delta-delta-CT method using the Applied Biosystems Step One Plus PCR System as we have previously described^72-77^. Primers used for qPCR are available upon request.

### Oral Glucose Tolerance Testing (OGTT)

The mice were fasted for 4 h before the tests. OGTT was performed after a single gavage of glucose, 2.5 g per kg body weight for standard diet and 1g per kg body weight for HFD and *ob/ob* studies. Blood glucose was then measured before (0 min) and after the injection (15, 30, 60, 120 min) using a OneTouch® SelectSimple™ glucometer (LifeScan Inc., China).

### Measurement of Plasma Hormone and Lipid Levels

Plasma insulin (EZRMI-13K, EMD Millipore) and corticosterone (501320, Cayman Chemical) levels were measured by ELISA. Plasma non-esterified fatty acid (HR Series NEFA-HR, Wako) and triglyceride (L-Type Triglyceride M, Wako) levels were measured using enzymatic assays according to manufacturer’s instructions.

### Indirect Calorimetry

To measure the effects of IMC on energy expenditure and physical activity mice were housed in metabolic cages (Oxymax CLAMS, Columbus Instruments) for indirect calorimetry measurements at room temperature (22°C). Mice were acclimated to the home cage system for 72 hours prior to data collection, and data were analyzed as previously described^72-74^.

### Cecal Microbiome Analyses

Snap frozen cecal DNA was isolated using the MO BIO Powersoil®-htp 96 well soil DNA isolation kit according to manufacturer’s instructions. Region-specific primers (515F/806R) were used for amplifying the V4 region of the bacterial 16S rRNA gene for high throughput sequencing using the Illumina HiSeq platform, paired end 150bp run. The reverse amplicon primer contains a 12-base Golay barcode sequence unique to each well that allows sample pooling for sequencing^78- 81^. More information can be found at the Earth Microbiome Project 16S rRNA Amplification Protocol where our protocols were adapted from: http://www.earthmicrobiome.org/emp-standard-protocols/16s/. Each sample was amplified in triplicate using 5 PRIME HotMaster Mix 2.5X (VWR 10847-708), combined, verified by 1.5% agarose gel and quantitated using Pico Green dsDNA Assay Kit (Thermofisher P7589). Samples were pooled (250 ng) and cleaned using the UltraClean PCR Clean-Up Kit protocol (Mo-Bio 12500-100). The quantified amplicons were sequenced with the Illumina HiSeq 2500 at the Broad Stem Cell Research Center at the University of California – Los Angeles on two lanes. The sequences were analyzed using the open source python software package Quantitative Insights Into Microbial Ecology (QIIME) version 1.9.1^78,83^ using default parameters for each step, except where specified. Demultiplexed sequences were aligned and clustered into operational taxonomic units (OTUs) based on their sequence similarity (97% identity) using the SortMeRNA/SumaClust open reference based OTU picking protocol in QIIME. Representative sequences for each OTU were aligned using PyNAST (a python-based implementation of NAST^84^ in Qiime and the Greengenes 11 database^85^. 38,826,537 total reads were generated after removal of singleton reads and rare (<0.01% of total reads) OTUs, with an average of 776,531 reads per sample. Samples were rarefied to the depth of the sample with the lowest number of reads (86,941sequences/sample) for beta diversity assessment only. Beta diversity was assessed using weighted UniFrac in QIIME. Adonis statistical test with 1,000 permutations was used to determine the strength and statistical significance of sample groupings. LEfSe was used with default parameters on OTU tables to determine taxa that best characterize each population^86^. Significant differences in relative abundance of taxa between groups and correlations with physiological parameters were assessed using the ALDEx2 package implemented in R^87,898^. Data were adjusted for false discovery rate using the Benjamini-Hochberg procedure and an adjusted p-value of p < 0.05 was considered statistically significant. All other plots were carried out using R (r-project.org).

### Untargeted Metabolomics

Moue plasma samples were prepared for untargeted metabolomics by diluting each plasma sample 1:20 in chilled methanol containing 5 internal standards as listed in the table below. The samples were then centrifuged at 14,000g for 20 minutes to precipitate out the protein pellet. The supernatant was recovered and subjected to LC-MS analysis. One-microliter aliquots taken from each sample were pooled and this QC standard was analyzed every 10 injection. The untargeted metabolomics was performed by injecting 7uL of each sample onto a 10 cm C18 column (Thermo Fisher CA) coupled to a Vanquish UHPLC running at 0.25mL/min using water and 0.1%formic acid as solvent A and acetonitrile and 0.1% formic acid as solvent B. The 15-min gradient used is given below. The Orbitrap Q Exactive HF was operated in positive and negative electrospray ionization modes in different LC-MS runs over a mass range of 50-750 Da using full MS at 120,000 resolution. Data dependent acquisitions were obtained on the pooled QC sample. The DDA acquisition (DDA) include MS full scans at a resolution of 120,000 and HCD MS/MS scans taken on the top 10 most abundant ions at a resolution of 30,000 with dynamic exclusion of 4.0 seconds and the apex trigger set at 2.0 to 4.0 seconds. The resolution of the MS2 scans were taken at a stepped NCE energy of 20.0, 30.0 and 45.0. XCMS was used to deconvolute the data using 5 ppm consecutive scan error, 5 to 60 seconds as minimum and maximum peak width, S/N threshold of 10, and span of 0.2 in positive mode and span of 0.4 in negative mode for retention time correction. The resulting peak table was further analyzed via MetaboLyzer. Briefly, the ion presence threshold was set at 0.7 in each study group. Data were then log-transformed and analyzed for statistical significance via non-parametric Mann-Whitney U-test (FDR corrected p-value <0.05). Ions present in just a subset of samples were analyzed as categorical variables for presence status via Fisher’s exact test. All p-values were corrected via the Benjamini-Hochberg step-up procedure for false discovery rate (FDR) correction. The data was then utilized for PCA, putative identification assignment, and pathway enrichment analysis via KEGG. In this dataset 7665 spectral features were detected, from which 1151 features were putatively assigned an identification in HMDB within a pre-defined 7ppm m/z error window. Also, the MS/MS spectra of 120 of these features matched with a score of greater than 50% to 120 unique compounds on the *mzCloud* database. Given the complicated nature of comparing the global metabolome across two treatment groups and 6 circadian time points, we used an algorithm called Selective Paired Ion Contrast Analysis (SPICA)^89^, to reveal subtle differences the plasma metabolome kinetically. Pairwise analysis was conducted between all adjacent timepoints (ZT2, ZT6, ZT10, ZT14, ZT18, and ZT22) for each treatment group (chow control versus chow + IMC), resulting in a total of 10 comparisons made (5 for chow and 5 for chow + IMC). Differences in the global plasma metabolome for each pairwise comparison were quantified via receiver operating characteristic (ROC) curve construction and area under the curve (AUC) calculations via Monte Carlo cross validation procedures in SPICA^89^. This analysis revealed that while all adjacent timepoints were roughly equally differentiated in the chow control data with an AUC averaging 0.884, this was not the case in the IMC-treated group. The AUC calculated for the IMC-treated group when comparing T10 vs. T14 was much greater than that of the chow-fed control group at the same time points (0.964 vs. 0.869), meaning the differences in the plasma metabolome between these two timepoints were much more pronounced in the IMC-treated mice compared to chow controls. Furthermore, The AUCs calculated for the subsequent two timepoint comparisons (T14 vs. T18 and T18 vs. T22) were much lower in the IMC-treated group (0.624 and 0.615, respectively) when compared to the chow control group (0.842 and 0.899, respectively), implying that the T14, T18, and T22 timepoints were poorly differentiated by IMC treatment. As a result, subsequent data analyses focused on this T10 to T14 transition, which also coincided with when the mice began to eat. The statistically significant positively-charged and negatively-charged spectral features which were present in more than 70% of the samples at ZT14 are shown in red in the volcano plots of Figure 4. These features were then putatively identified in the Human Metabolome and the KEGG databases using their accurate mass-to-charge (m/z) values within a 7ppm error window. The KEGG annotated pathways associated with these putative metabolites were then identified. Figure 4c represents such KEGG pathways associated with the negatively charged spectral features in this study. This figure displays the KEGG metabolic pathways with the highest statistically significance to which the ions were assigned. The blue and red bars are the unadjusted and the FDR (false discovery rate)-adjusted -log of p-values respectively, while the orange line marks the significance threshold. This figure shows that lipid metabolism is the most perturbed metabolic pathway at ZT14.

### Data Analyses for Circadian Rhymicity (Cosinor Analyses)

A single cosinor analysis was performed as previously described^90-92^. Briefly, a cosinor analysis was performed on each sample using the equation for cosinor fit as follows:

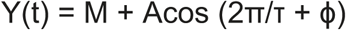

where M is the MESOR (Midline Statistic of Rhythm, a rhythm adjusted mean), A is the amplitude (a measure of half the extent of the variation within the cycle), ϕ is the acrophase (a measure of the time of overall highest value), and τ is the period. The fit of the model was determined by the residuals of the fitted wave. After a single cosinor fit for all samples, linearized parameters were then averaged across all samples allowing for calculation of delineralized parameters for the population mean. A 24-h period was used for all analysis. Comparison of population MESOR, amplitude, and acrophase were performed as previously described^28^. Comparisons are based on F-ratios with degrees of freedom representing the number of populations and total number of subjects. All analysis was done in R v4.0.2 using the cosinor and cosinor2 packages^93-95^.

### Statistical Analysis

All data were analyzed using either one-way or two-way analysis of variance (ANOVA) where appropriate, followed by either a Tukey’s or Student’s t tests for post hoc analysis. Differences were considered significant at p <0.05. All mouse data analyses were performed using Graphpad Prism 6 (La Jolla, CA) software.

## ACKNOWLEDGEMENTS

This work was supported by National Institutes of Health grants R01 HL120679 (J.M.B.), P01 HL147823 (J.M.B., S.L.H.), P50 AA024333 (J.M.B), U01 AA026938 (J.M.B.), P50 CA150964 (J.M.B.), R01 HL103866 (S.L.H.), R01 HL147883 (A.J.L.), R01 HL144651 (A.J.L. and Z.W.), R01 HL130819 (Z.W.), F32 DK122623 (C.M.G.), T32 DK007307 (C.M.G.), a Leducq Transatlantic Networks of Excellence Award (S.L.H.), and the American Heart Association (Postdoctoral Fellowships 17POST3285000 to R.N.H and 15POST2535000 to R.C.S). Development of some of the mass spectrometry methods reported here were supported by generous pilot grants from the Clinical and Translational Science Collaborative of Cleveland (4UL1TR000439) from the National Center for Advancing Translational Sciences (NCATS) component of NIH and the NIH Roadmap for Medical Research, the Case Comprehensive Cancer Center (P30 CA043703), the VeloSano Foundation, and a Cleveland Clinic Research Center of Excellence Award.

## AUTHOR CONTRIBUTIONS

R.C.S., C.M.G., and J.M.B. planned the project, designed experiments, analyzed data, and wrote the manuscript; A.K., J.C.G-G., A.J.L., and S.L.H. help design experiments and provided useful discussion directing collaborative aspects of the project; R.C.S., C.M.G., R.N.H., A.L.B., A.B., C.F., K.K.F., F.M.A., D.F., A.D.G., C.N., A.M., J.A.B., J.T.A., M.M., M.G., B.W., T.D.M., A.R.A., G.S., A.K., A.J.L., and J.M.B. either conducted mouse experiments, performed biochemical workup of mouse tissues, analyzed data, and aided in manuscript preparation; All authors were involved in the editing of the final manuscript.

## Abbreviations used

BMAL1: aryl hydrocarbon receptor nuclear translocator like
*Cry1*: cryptochrome 1
CutC: choline trimethylamine lyase
CVD: cardiovascular disease
DIO: diet-induced obesity
*Fmo3*: flavin containing monooxygenase 3
IMC: iodomethylcholine
LPC: lysophosphatidylcholine
*Per2*: period 2
PC: phosphatidylcholine
PCA: principal component analysis
RevErb*α*: nuclear receptor subfamily 1 group D member 1
TMA: trimethylamine
TMAO: trimethylamine N-oxide
ZT: zeitgeber.

## FIGURE LEGENDS

**Figure 1 – figure supplement 1.**
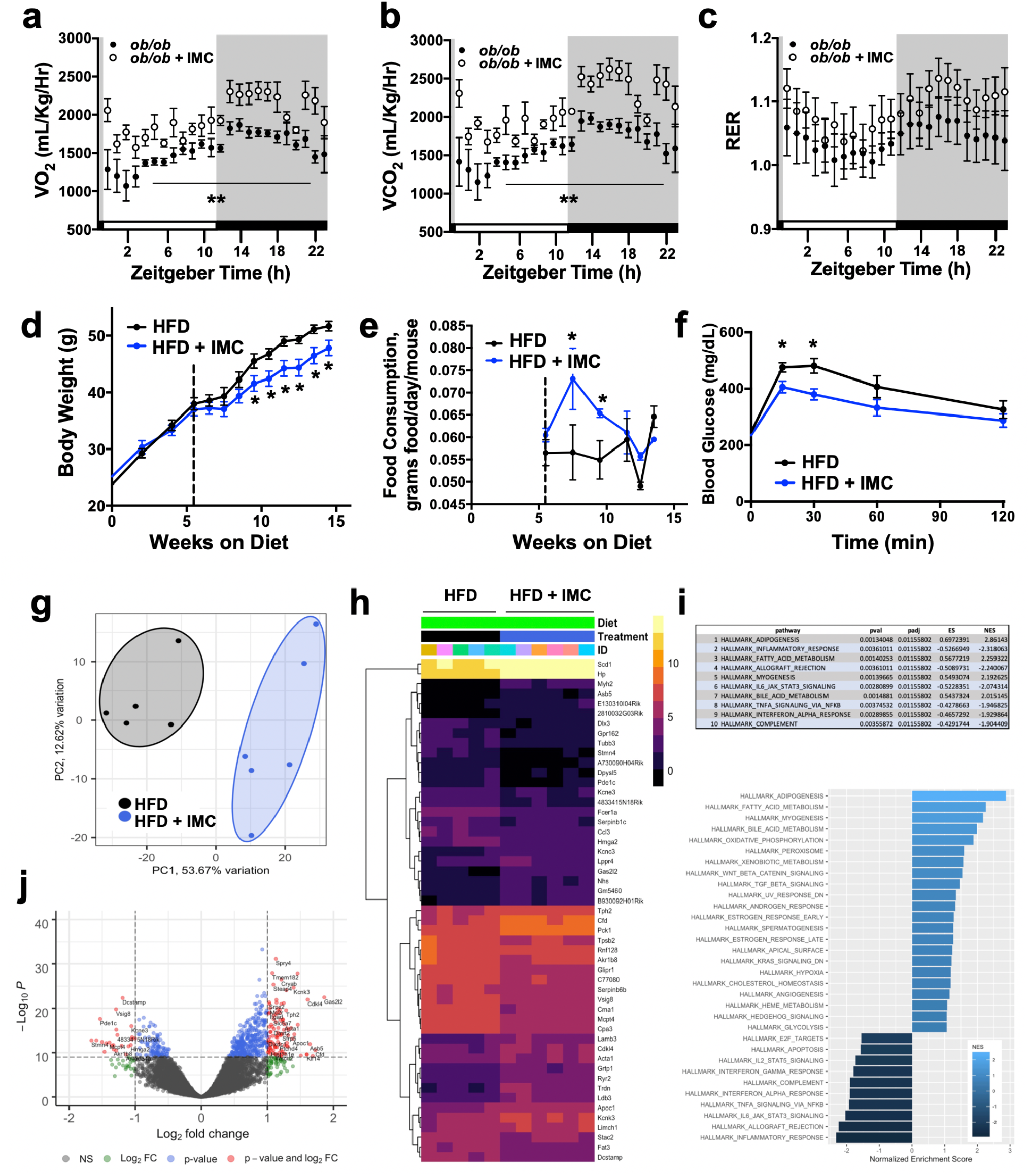
Choline TMA Lyase Inhibition Increases Energy Expenditure and Alters Gene Expression in White Adipose Tissue. *Related to main text Figure 1*. Panels **a**-**c** represent indirect calorimetry data from control and IMC-treated *ob/ob* mice. **a**. oxygen consumption. **b**. CO_2_ production. **c**. Respiratory exchange ratio collected in the Oxymax CLAMS metabolic cage system. **d-f**. C57BL6 mice fed a high fat diet (HFD) for 6 weeks to establish obesity (body weight > 35 grams); after 6 weeks of DIO mice were continued on HFD alone of HFD containing IMC for another 10 weeks to test whether IMC can improve obesity-related phenotypes. **d**. body weight curve. **e**. food consumption after drug administration. **f**. glucose tolerance testing 8 weeks following IMC administration. **g-k**. Male C57BL6/J mice were fed a HFD for 20 weeks and then gonadal white adipose tissue was collected for examination of global transcriptome alterations via RNA sequencing. **g**. Principal component analysis. **h**. heatmap of most significantly altered mRNA transcripts. **i**. Hallmark pathway analysis of differentially expressed genes (DEGs). **j**. volcano plot showing most significantly up and down regulated transcripts.

**Figure 2 – figure supplement 1.**
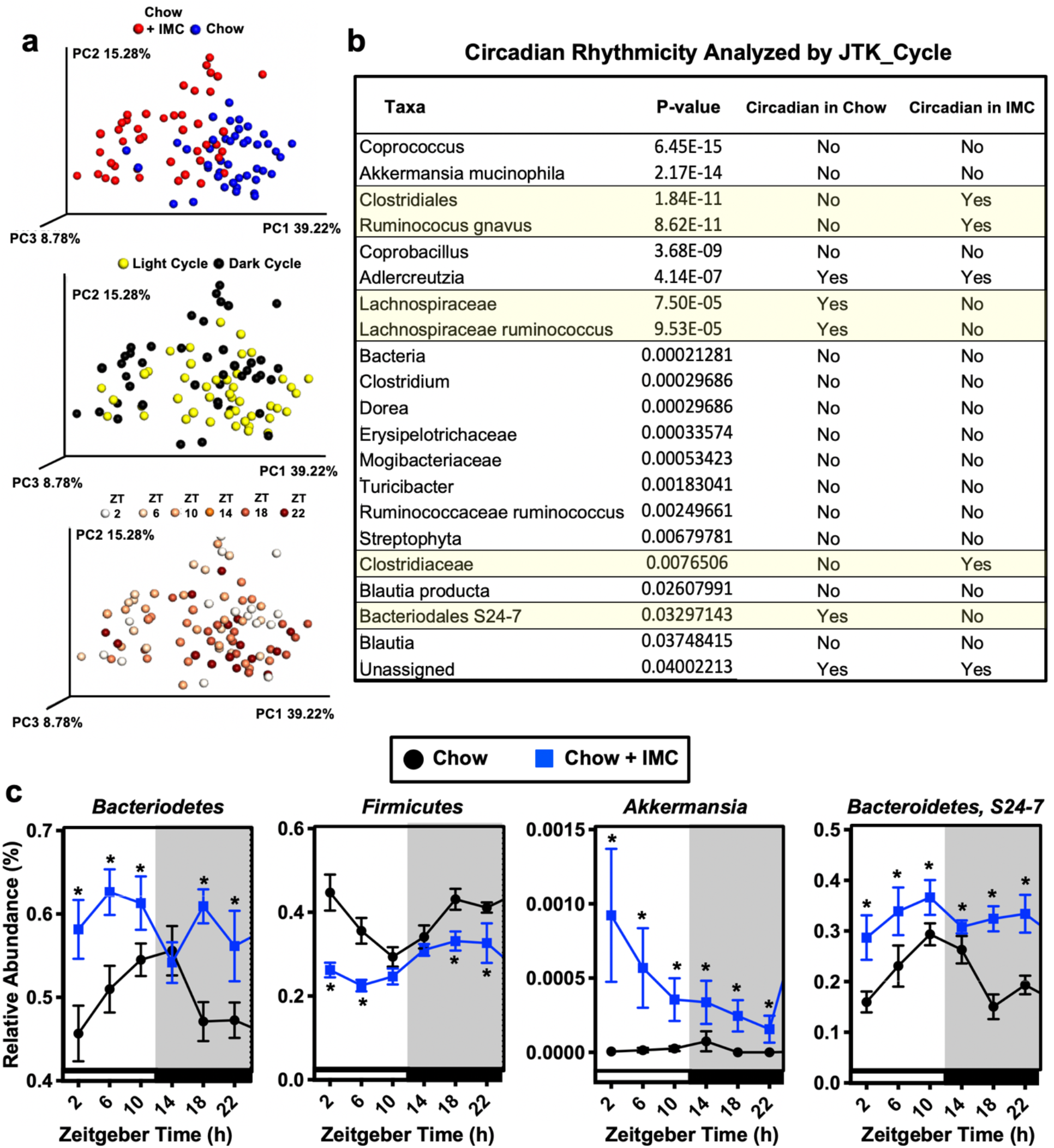
TMA Lyase Inhibition Beneficially Improves Circadian Oscillations in Gut Microbial Communities. *Related to main text Figure 2*. Wild type male C57BL/6J mice were fed chow or chow supplemented with the choline TMA lyase inhibitor iodomethylcholine (IMC) for 7 days. Mice were then necropsied at 4 hour intervals to collect cecum for 16S-based microbiome analyses. **a**. Principal coordinate analysis (Bray-Curtis) comparing chow versus chow + IMC, light cycle versus dark cycle, and across all ZT time points. **b**. Identification of cyclic taxa whose relative abundance also differed significantly between groups (highlighted in yellow). FDR corrected P values for treatment effect (two-way ANOVA) are shown. Circadian rhythmicity of cecal microbial taxa was analyzed using JTK_Cycle. **c**. Relative abundance of key IMC-altered taxa that are circadian in nature (n = 7-8); Significance between groups at specified zeitgeber (ZT) time points were compared using Student’s t-test (p <0.05).

**Figure 3 – figure supplement 1.**
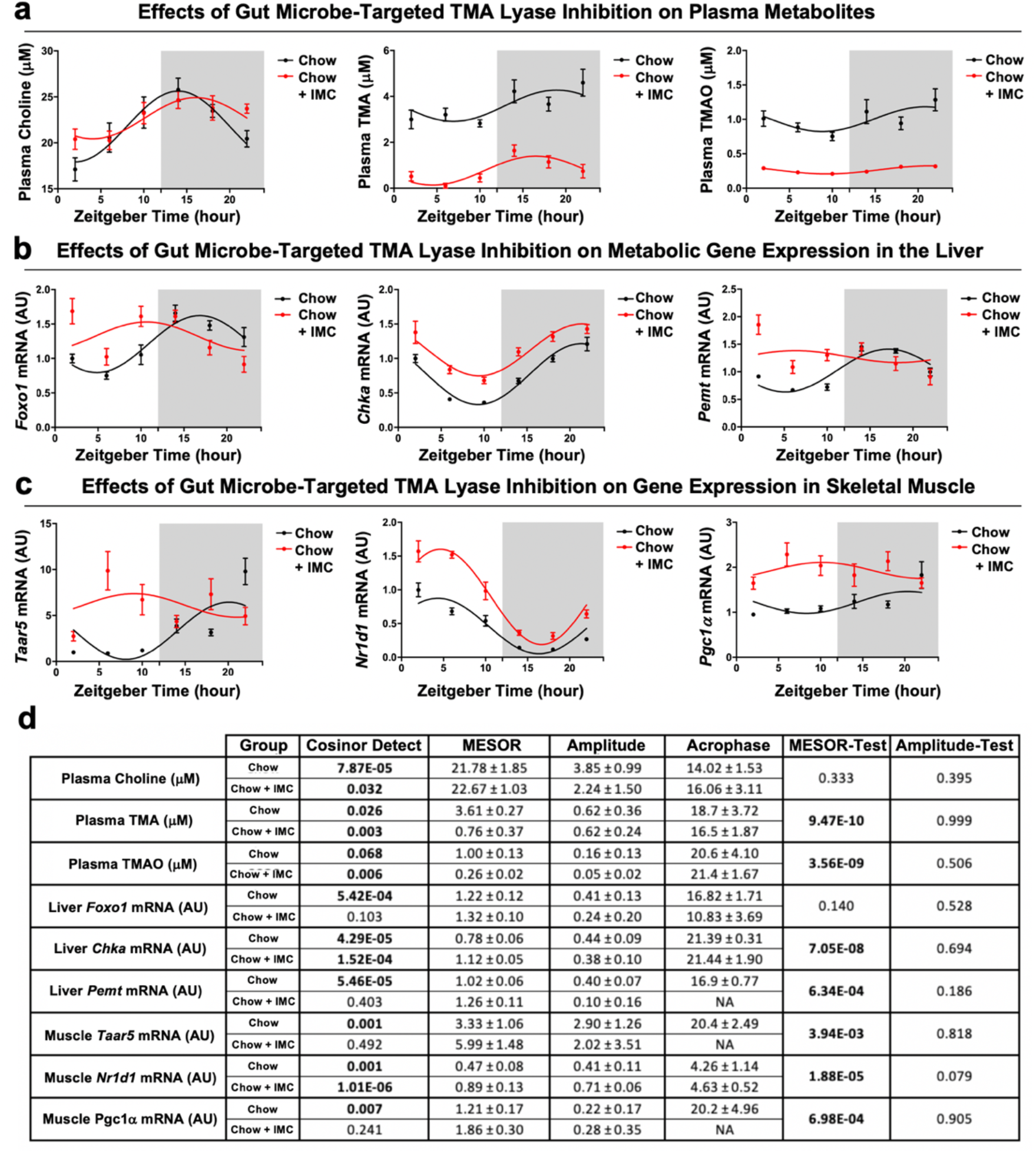
TMA Lyase Inhibition Alters the Circadian Rhythmicity of Host Metabolism-Associated Gene Expression. *Related to main text Figure 3*. Wild type male C57BL/6J mice were fed chow or chow supplemented with the choline TMA lyase inhibitor iodomethylcholine (IMC) for 7 days. Mice were then necropsied at 4 hour intervals to collect blood and metabolic tissues (liver and gastrocnemius skeletal muscle). **a**. Plasma levels of choline, trimethylamine (TMA), and trimethylamine N-oxide (TMAO). **b**. The relative mRNA expression of forkhead box protein O1 (*Foxo1*), choline kinase *α* (Chk*α*), phosphatidylethanolamine N-methyltransferase (*Pemt*) in the liver over a 24-hour period. **c**. The relative mRNA expression of the TMA receptor trace amine-associated receptor 5 (*Taar5*), nuclear receptor subfamily 1 group D member 1 (*Nr1d1*; RevErb*α*), and peroxisome proliferator-activated receptor gamma coactivator 1 *α* (Pgc1*α*) in the gastrocnemius muscle over a 24-hour period. **d**. Circadian parameters (MESOR, amplitude, and acrophase) were determined by cosinor analysis with a 24 hour period in plasma, liver, and skeletal muscle. The overall fit for each model is shown in Cosinor Detect, and a Wald multivariate test was used to compare circadian parameters between groups. All data represent mean -/+ S.E.M (n = 7-8 per group).

**Figure 4 – figure supplement 1.**
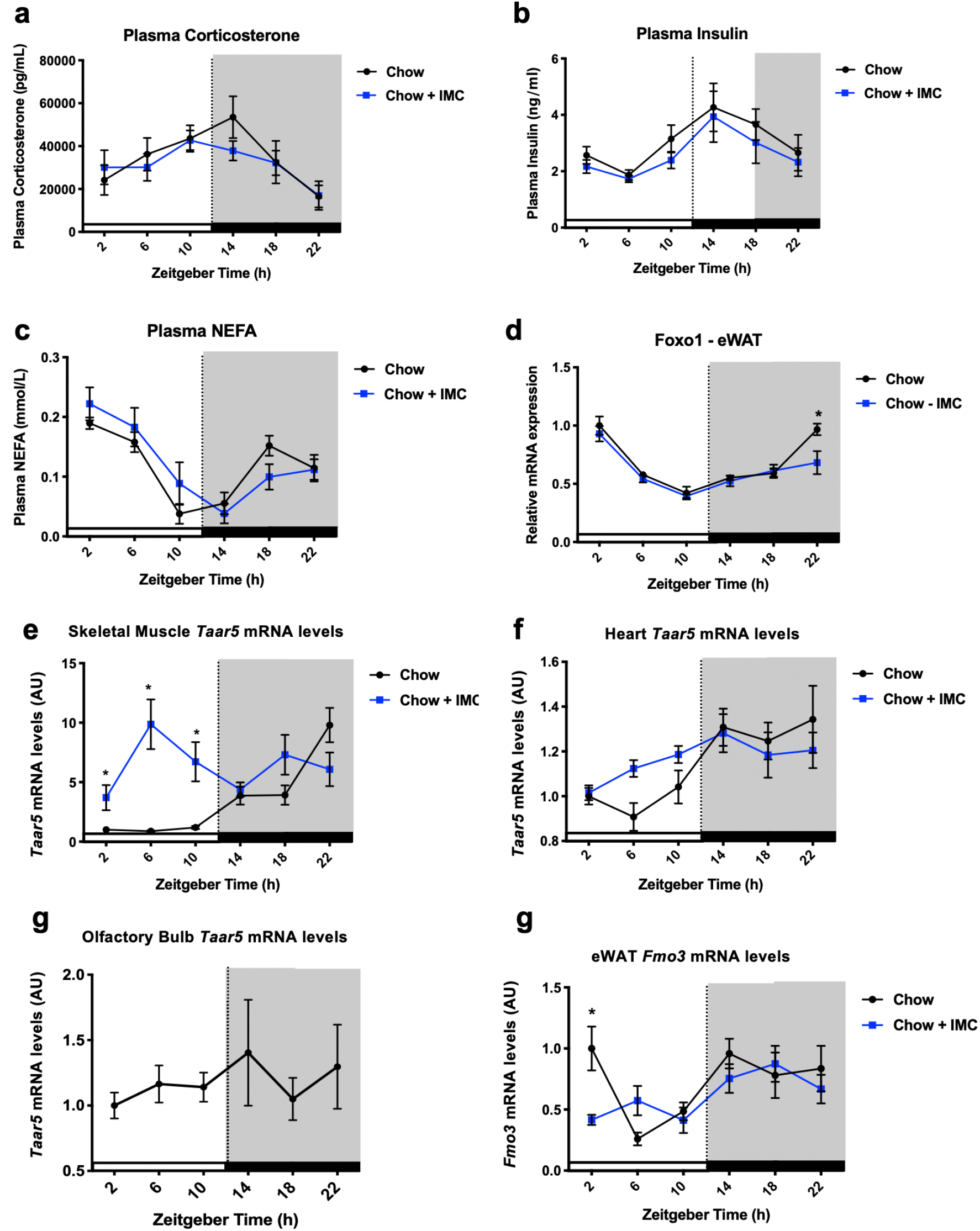
Choline TMA Lyase Inhibition Impact on the Circadian Rhythmicity of Host Hormone Levels and Gene Expression. *Related to main text Figure 4*. Wild type male C57BL/6J mice were fed chow or chow supplemented with the choline TMA lyase inhibitor iodomethylcholine (IMC) for 7 days. Mice were then necropsied at 4 hour intervals to collect blood/plasma. **a**. Plasma corticosterone levels were measured via ELISA. **b**. Plasma insulin levels were measured by ELISA. **c**. Plasma non-esterified fatty acid (NEFA) levels were measure via enzymatic assay. **d**. *FoxO1* mRNA levels in epidydimal white adipose tissue (eWAT). **e**. *Taar5* mRNA levels in gastrocnemius skeletal muscle. **f**. *Taar5* mRNA levels in the heart. **g**. *Taar5* mRNA levels in the olfactory bulb in chow-fed mice. **h**. *Fmo3* mRNA level in epidydmal white adipose tissue (eWAT). Significance between groups (n = 7-8) at each zeitgeber time point were compared using Student’s t-test (p <0.05).

